# The contribution of eye movements to memory retrieval depends on the visual input

**DOI:** 10.1101/2023.02.19.529117

**Authors:** Keren Taub, Shlomit Yuval-Greenberg

**Affiliations:** Sagol school of neuroscience, Tel-Aviv University; School of psychological sciences, Tel-Aviv University

## Abstract

When attempting to recall previously seen visual information, people often move their eyes to the same locations where they initially viewed it. These eye-movements are thought to serve a role in enhancing memory retrieval, although the exact mechanism underlying this effect is yet unknown. To investigate this link between eye-movements and memory, we conducted an experiment with 80 adult participants. Participants were asked to perform a memory retrieval task, while viewing either the same visual context as during encoding or an altered one.

Results showed that the benefit of eye movements to memory retrieval was dependent on the visual input. This suggests that the contribution of eye-movements to memory may not be from the motor behavior itself, but from its visual consequences. Our findings thus challenge the hypothesis that eye movements act as a motor retrieval cue and support the view that their visual consequences act as a sensory one.

**Statement of Relevance:** An intriguing question in cognition is how humans encode memorized material and what helps them retrieve it. It is known that when an action or stimulus is repeated both when information is encoded and when it is retrieved, this can act as a ‘retrieval cue’ and enhance memory performance. It is also known that people tend to reenact the same eye movements during retrieval as they did during encoding, and this behavior is associated with higher memory performance. This has led to the hypothesis that eye movements act as a retrieval cue. However, we challenge this hypothesis by showing that the visual *consequences* of eye movements, rather than the motor action that accompanies them, is the key factor for memory enhancement. Understanding the factors that influence memory provides crucial insight into the relationship between external behaviors and internal memory processes, leading to significant implications for the educational and clinical settings.

## Introduction

The main purpose of eye movements is to guide vision, moving the most sensitive part of the retina (the *fovea*) to objects of interest to enhance visual acuity. However, evidence shows that during cognitive tasks, people often move their eyes to empty locations in space where no objects of interest are present (Abeles & Yuval-Greenberg, 2017; Orquin & Mueller Loose, 2013; Renkewitz & Jahn, 2010; Salvi & Bowden, 2016). Specifically, when trying to remember visual information, people tend to move their eyes to the locations where this information was previously presented, even when it no longer does. Research further shows that during memory retrieval, people tend to reenact the same eye movements they have performed while encoding the information (Altmann, 2004; Brandt & Stark, 1997; Henderson & Castelhano, 2005; Hoover & Richardson, 2008; Irwin, 1992; Scholz et al., 2016; Spivey & Geng, 2001; Zelinsky & Loschky, 2005).

Studies have suggested that the reenactment of eye movements and the tendency to re-fixate the locations where targets were previously presented during retrieval serves a functional role in enhancing memory performance (Laeng et al., 2014; Rosner et al., 2022; Scholz et al., 2016, 2018). This view was supported by a study by Johansson & Johansson (2014), in which participants were asked to memorize the orientation and spatial location of objects and were later tested on them. When allowed to free-view the screen during test, participants tended to gaze more often at the screen quadrant where the target object was originally presented. When participants were not allowed to free-view but were asked to fixate on specific locations during the test, their performance depended on the location of fixation – recall performance was higher when the fixated location was congruent with the location where the recalled object was previously presented. These results indicate that gazing at the location where a retrieved object was previously presented, is beneficial for memory performance compared to gazing at different locations, supporting the hypothesis that eye movements enhance memory retrieval.

Despite strong evidence on the role of eye movement in memory enhancement, the mechanism underlying this effect is poorly understood, and it remains hitherto unknown what it is about eye movements that causes this enhancement. One possible explanation is that the oculomotor behavior is associated with a stored memory and its execution helps resurface the relevant memory. This “motor hypothesis” is consistent with studies of other, non-oculomotor, behaviors that showed how bodily motions performed during encoding can enhance memory performance when reenacted during retrieval (Dijkstra et al., 2007).

However, eye movements are not a purely-motor mechanism, as each eye movement results in a direct and immediate sensory consequence – the shifting of the retinal image. During retrieval, when gaze is shifted to the same location where it was located during encoding, much of the visual context is recreated. Although the memorized item itself is no longer present during retrieval, the rest of the peripheral context is available and could serve as a visual retrieval cue. This “visual hypothesis” suggests that eye movements are linked to memory enhancement, not because they constitute a motor retrieval cue, but because their *consequence* is a sensory retrieval cue. Here we examine this hypothesis relative to the alternative hypothesis that eye movements act as a motor retrieval cue.

The role of the visual context in memory retrieval has not yet been fully explained. One previous study examined the effect of the visual context on the reenactment of eye movements during memory retrieval (Johansson et al., 2006). In that study, retrieval took place in conditions of complete darkness, eliminating the entire visual context. The finding showed that even in complete darkness, eye movements performed during encoding were reenacted during retrieval. This could suggest that the link between eye movements and memory retrieval does not dependent on the visual context. However, while eye movements performed during retrieval were found to be similar to those performed during encoding, even in darkness, their functional role in memory performance remained unknown. In other words, it was shown that reenactment of eye movements exists even without visual input, whether or not it plays a role in *enhancing* memory performance remains hitherto unknown. This last step is crucial to support or refute the hypothesis that eye movements act as a retrieval cue.

Here we examine the hypothesis that the functional role of eye movements in memory retrieval depends on the presence of visual context. Using a similar design to that used by Johansson and Johansson (2014), we examined the effect of visual context on the link between gaze behavior during memory retrieval and memory performance. Participants were presented with objects on four screen quadrants and were asked to memorize their orientation and location in relation to other objects. During test, the visual input was either similar to encoding, preserving the same visual context, or altered by a physical occluder. While both settings required participants to retrieve visual information that was no longer available, if memory-related eye movements rely on the reinstatement of visual information on the retina – the alteration of the visual context would eliminate any beneficial effect stemming from the reenactment of similar eye movements. In contrast, if memory-driven eye movements rely on motor retrieval cues – alteration of the visual context would not affect the benefit of eye movements. Through this design, we were able to examine the isolated effect of visual input on the functional role of eye movement in memory retrieval. The findings reveal that the beneficial effect of eye movements on memory performance depends on the visual context. These findings challenge the hypothesis that eye movements act as a motor retrieval cue and support the alternative hypothesis that their visual consequence act as a sensory one.

## Methods

### Participants

A total of 80 individuals participated in the experiment in two groups of 40 each (informed group: 31 females, age range 21-30, Mean age = 23.80, SD age = 2.027; Naïve group: 28 females, age range 18-40, Mean age = 24.775, SD age = 3.l725). Sample size was chosen based on power analysis using Gpower 3.1 (Faul et al., 2007), with an estimation of a medium effect size (Cohen’s d=0.5), significance level of 0.05 and power of 0.8. Power estimation indicated a required sample size of at least N=34 for each group. Participants were recruited through an online recruitment system, and received course credit for their participation. All participants had normal or corrected-to-normal vision and were native Hebrew speakers. Participants reported having no known psychiatric or neurological deficits. Participants signed an informed consent prior to the experiment. The experiment was approved by the ethics committee of Tel Aviv University and the School of Psychological Sciences.

### Stimuli

Stimuli consisted of 96 visual objects, chosen from Snodgrass and Vanderwart’s object pictorial set (Snodgrass & Vanderwart, 1980) and Multipic database (Duñabeitia et al., 2018; http://www.bcbl.eu/databases/multipic). The objects were divided into 16 groups of six objects each, according to category (e.g vehicles, animals, etc.). Objects for this experiment were selected based on a preliminary research in which 200 objects were presented to 20 native Hebrew participants who were asked to indicate the name and orientation of each object. Only objects whose identity and orientation were widely agreed upon by at least 90% of participants, were included in the main experiment.

### Procedure

Participants were divided to two groups: informed and naïve. Prior to the beginning of the experiment, participants of the informed group were told that research suggests that looking at the locations where items were previously presented, helps remembering them. These participants were further reminded of this information prior to each test block and were encouraged to adopt this strategy to improve their performance. In contrast, participants of the naïve group were not told this information and were not encouraged to adopt a specific strategy.

The experiment included four blocks, each starting with an encoding phase (3 minutes), followed by a test phase (approximately 7 minutes). The encoding phase of each block was identical to that used by Johansson & Johansson (Johansson & Johansson, 2014). The screen was divided into four quadrants by two crossing black lines. A group of six objects of one category was presented at random locations in one quadrant for 30 seconds and then replaced by the next group on another quadrant. In each group of objects, half of the objects (randomly picked) were oriented to the right and half to the left. This was repeated for all four quadrants, with their order randomly picked. After this, all objects were presented again at the same locations as before but this time together for another 60 seconds. Participants were asked to memorize objects’ location and orientation.

At each trial of the test phase, participants listened to pre-recorded statements describing either an object’s orientation or its relative location, similar to Johansson & Johansson (2014). Participants were instructed to respond by pressing one of six keys, indicating both their response to the statement (True / False) and their confidence level (from 1 – weakest, to 3 – strongest, with 1 defined as a complete guess). Confidence ratings were not included in the analysis of the present study. Before the test phase of each block, an experimenter entered the room. In two of the four blocks (first and third or second and fourth, counterbalanced among participants), the experimenter placed a black cardboard in front of the screen, occluding most of the visual scene from view (No-visual context condition). In the other two blocks (second and fourth or first and third, correspondingly), they left the room immediately after telling participant that the test phase is about to begin, but without altering anything in the room setup (Visual context condition). In the informed group, the experimenter reminded the participant before each memory test that looking at the locations where items were previously presented could enhance performance. At the end of each block participants received feedback on their performance with the percent of correct responses.

To ensure that participants recognized the objects’ names and orientations (as determined by the preliminary study), they were presented with all of the objects at the end of the experiment and were asked to name them and report their orientations. Objects whose names or orientations were not correctly recognized by a specific participant were excluded from analysis for that participant, with an average of 2.13% [SD=2.07%] of the objects excluded per participant.

### Apparatus and Eye tracking

Participants were seated in a sound-attenuated room at a distance of 100 cm from a 24 inch ASUS VG248QE LCD screen, with their head rested on a chin-rest. Binocular eye-movements were monitored using a remote infrared video-oculographic system (EyeLink 1000 Plus, SR Research Ltd, Ontario, Canada), with a sampling-rate of 1000 Hz, spatial resolution <0.01° and average accuracy of 0.25°-0.5° when using a head-rest, as reported by the manufacturer. At the beginning of each block, participants completed a 9-point eye tracking calibration procedure performed with the same illumination conditions as the main experiment. Calibration was accepted when validation confirmed that fixation errors were of no more than 0.2°.

### Analysis

Analysis focused on trials of test phase, according to group and condition. The first aim was to examine for how long participants gazed at the *congruent* quadrant, the quadrant in which the target object was previously presented, relative to the other quadrants (the *incongruent* quadrants, numbered 1, 2, and 3 in a clockwise order relative to the congruent quadrant). To this aim, gaze dwell time proportion was calculated per quadrant and trial as the total fixation time during which participants gazed at this quadrant relative to the total time of the trial.

Fixations were considered valid and included in the analysis if they were longer than 100 ms and were located within the borders of the screen (even in the No-visual context condition in which the screen was invisible). Trials with no fixations within screen limits were omitted from analysis (mean percent of trials included in the analysis across participants: 94.694%, SD: 2.948%, range 74.48%-99.48%). For each participant we calculated the average dwell time proportion of the congruent quadrant by averaging across trials, and the average dwell time proportion of the incongruent quadrants, by averaging across trials *and* across the three incongruent quadrants (defined differently for different trials). Next, we examined the link between dwell time congruity and memory performance. This was done by first detecting the trials in which dwell time on the congruent quadrant - the quadrant where the target object was previously located - was higher than chance (25% of the time). Accuracy rates on these trials, defined as congruent-gaze trials, were compared with accuracy on the rest of the trials, defined as incongruent-gaze trials. All eye movements analysis was performed using Matlab R2021b.

## Results

### Accuracy rates

The average accuracy rate across conditions was 83.75% (SD=7.79%, range: 65.38%- 97.81%). We conducted a 2×2 ANOVA with between variable of Group (naïve / informed) and within variable of Context (visual context / no-visual context). There was no evidence for a main effect of Group or Context, or for an interaction between the two (Group effect: F(1)=2.202, p=0.142; Context effect: F(1)=3.039, p=0.085, marginal significance indicating a slightly better performance in the no-visual context than the visual context condition; Interaction effect: F(1)<1, p=0.799 ;). These null results where further supported by Bayesian statistics Group effect: BF_01_=1.332, % error=1.267; Context effect: BF_01_=1.472, % error=2.196; Interaction effect: BF_01_=8.138, % error=3.125;).

### Dwell time analysis: where do people look when they try to remember?

We performed a 2×2×2 ANOVA, with three independent variables: Group (naïve / informed), Context (visual context / no-visual context) and Quadrant congruency (congruent / incongruent). The first variable was measured between-participants, and the other two within-participants. The dependent measurement was the average dwell time proportion – the proportion of time in which gaze was directed towards the quadrants. This measurement was calculated for each participant and condition and separately for congruent and incongruent quadrants, with the incongruent quadrant representing the average across the three incongruent quadrants. Consistently with previous studies, we found a significant main effect of Quadrant congruency (F(1)=182.991, p<0.001), supporting previous evidence suggesting that gaze tends to be directed more towards the congruent quadrant than to the incongruent quadrants.

The main purpose of this study was to examine how visual context modulates the congruency effect. Interestingly, we found that the effect was evident not only when the visual context was present (t(107.54)=14.61, p<0.001), as was found by previous studies, but also when it was absent (t(107.54)=10.12, p<0.001). In other words, participants tended to gaze at congruent locations, even when the visual context was occluded. A significant two-way interaction between Context and Quadrant congruency revealed that, although the effect occurred both with and without visual context, it was *larger* when the visual context was present, relative to when it was absent (F(1)=30.675, p<0.001) (see Figure 1).

**Figure 1.**
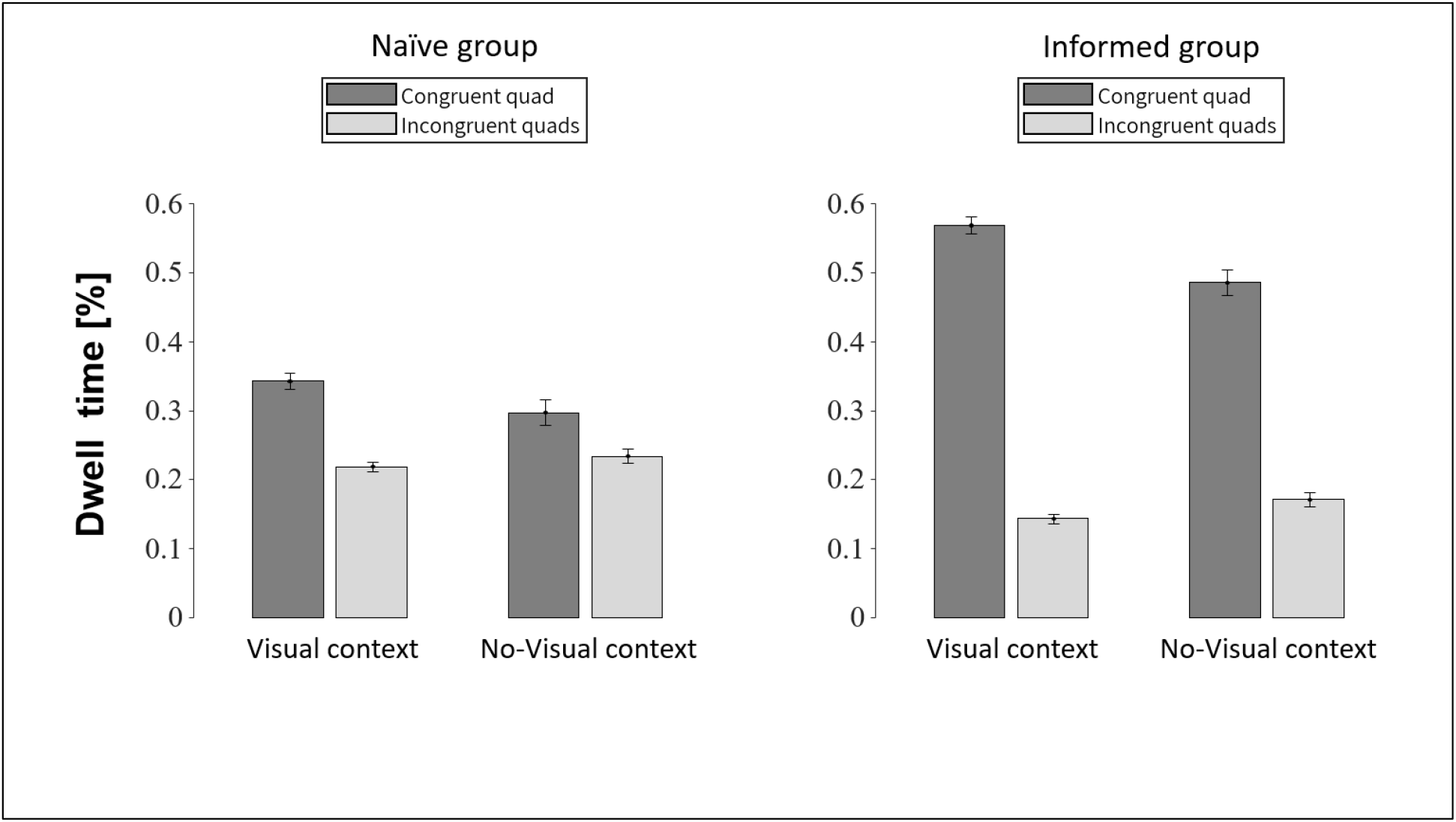
Dwell time on congruent and incongruent quadrants. Participants gazed more at the congruent than the incongruent quadrant in both conditions and groups.

Another question was whether informing participants explicitly on the advantages of gazing at location where memorized items have previously appeared would affect their behavior and improve their performances. To this aim we have examined the two-way interaction between Quadrant congruency and Group and found it to be significant (F(1)=65.971, p<0.001), indicating a larger congruity effect for the informed participants compared with the naïve participants. Simple effect analyses performed separately on the two groups suggested that participants of both groups tended to gaze more towards congruent rather than incongruent quadrants (naive group: t(78)=3.82, p<0.001; informed group: t(78)=15.31, p<0.001) (see Figure 1).

Simple effect analysis revealed a significant difference between dwell time for congruent and incongruent quadrant for each group and condition (Naïve group, visual context condition: t(58.27)=6.294, p<0.001; Naïve group, no-visual context condition: t(58.27)=3.192, p<0.01; Informed group, visual context condition: t(52.122)=13.276 p<0.001; Informed group, no-visual context condition: t(52.122)=9.909, p<0.001). This indicates that the tendency to gaze more at the congruent quadrant was evident both with and without visual context and in participants who were informed as well as in those who were naïve. These findings are depicted in Figure 1 and Figure 2.

**Figure 2.**
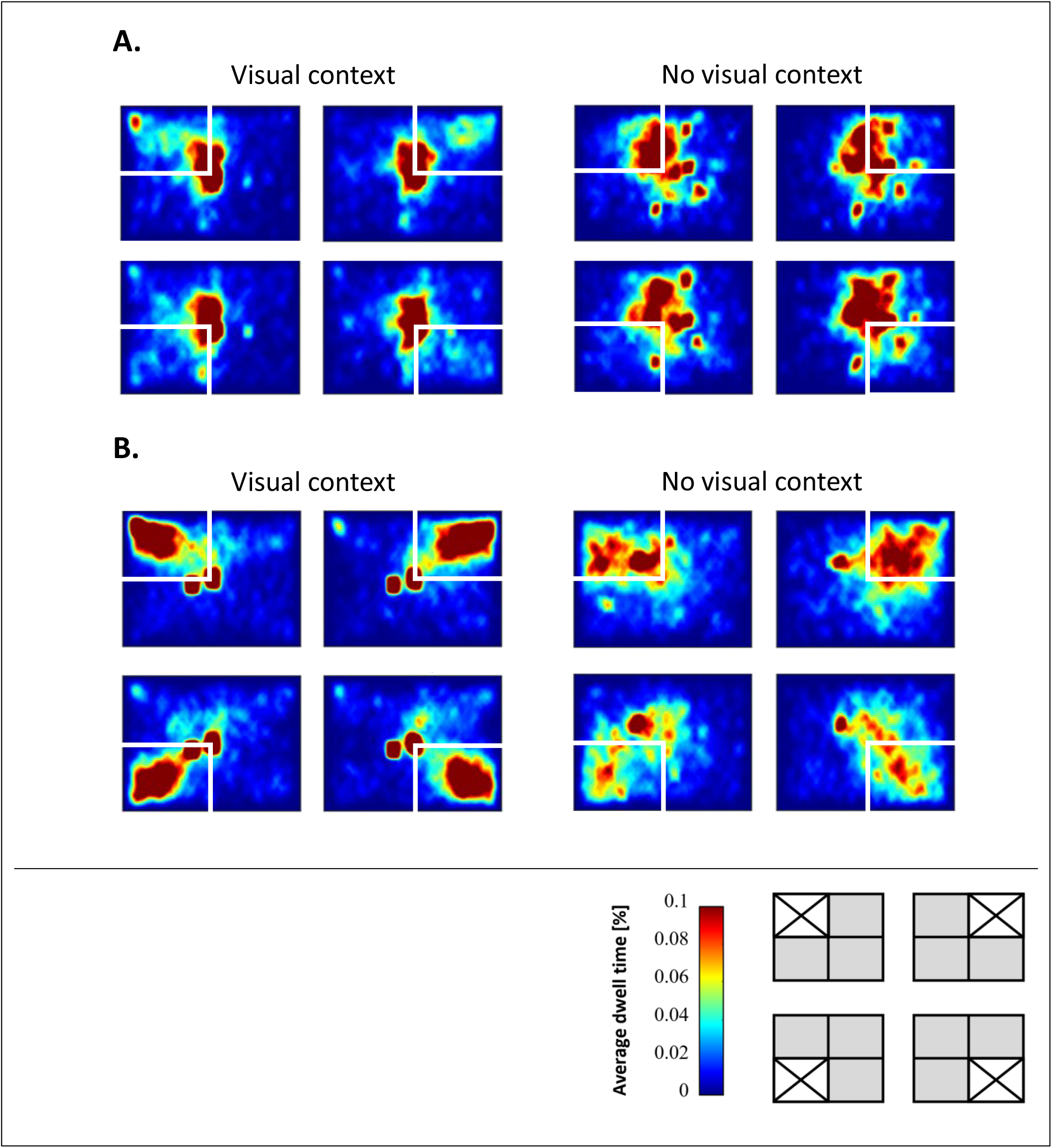
Gaze distribution across the visual field, according to group and visual context (A. naïve group, B. informed group). All graphs are divided into four panels, each describes gaze distribution according to target location: each of the four quadrants. The panel here on the right is an example for the division of the four panels, with the white area indicating the location of the target objects.

Finally, we found no evidence for difference between the groups in the way the congruity effect was modulated by visual context. This was evident by an insignificant three-way interaction (F(1)=2.409, p=0.125).

### The link between dwell time and accuracy rates: how does looking at congruent locations affects memory performance?

Next, we examined whether the tendency of participants to gaze at congruent locations enhanced their memory performance, and whether this enhancement was linked to the presence of a visual context. For this analysis, test trials were divided to congruent test trials (trials in which the average dwell time proportion for the congruent quadrant was more than 25%) and incongruent test trials (the rest of the trials).

We conducted a 2×2×2 ANOVA with three independent variables: Group (naïve / informed), measured between-participants, and Context (visual context / no-visual context) and Gaze congruency (congruent / incongruent), measured within-participants.

This analysis revealed a significant main effect for Context, showing higher accuracy rates for the no-visual context compared with visual context condition (F(1)=6.047, p=0.016).

Analysis further revealed a main effect of Gaze congruency, with congruent gaze associated with higher accuracy rates than incongruent gaze (F(1)=4.214, p=0.0436). However, this effect was modulated by the visual context, as suggested by a significant two-way interaction between Gaze congruency and Context (F(1)=6.402, p<0.05 (p=0.0135). Simple effect analysis for this interaction revealed that in the visual context condition, but not in the no-visual context condition, there was a significant difference between congruent and incongruent gaze (visual context condition: t(147.84)=3.246, p=0.001; no-visual context condition: t(147.84)=-0.391, p=0.697). Figure 3 depicts these findings.

**Figure 3.**
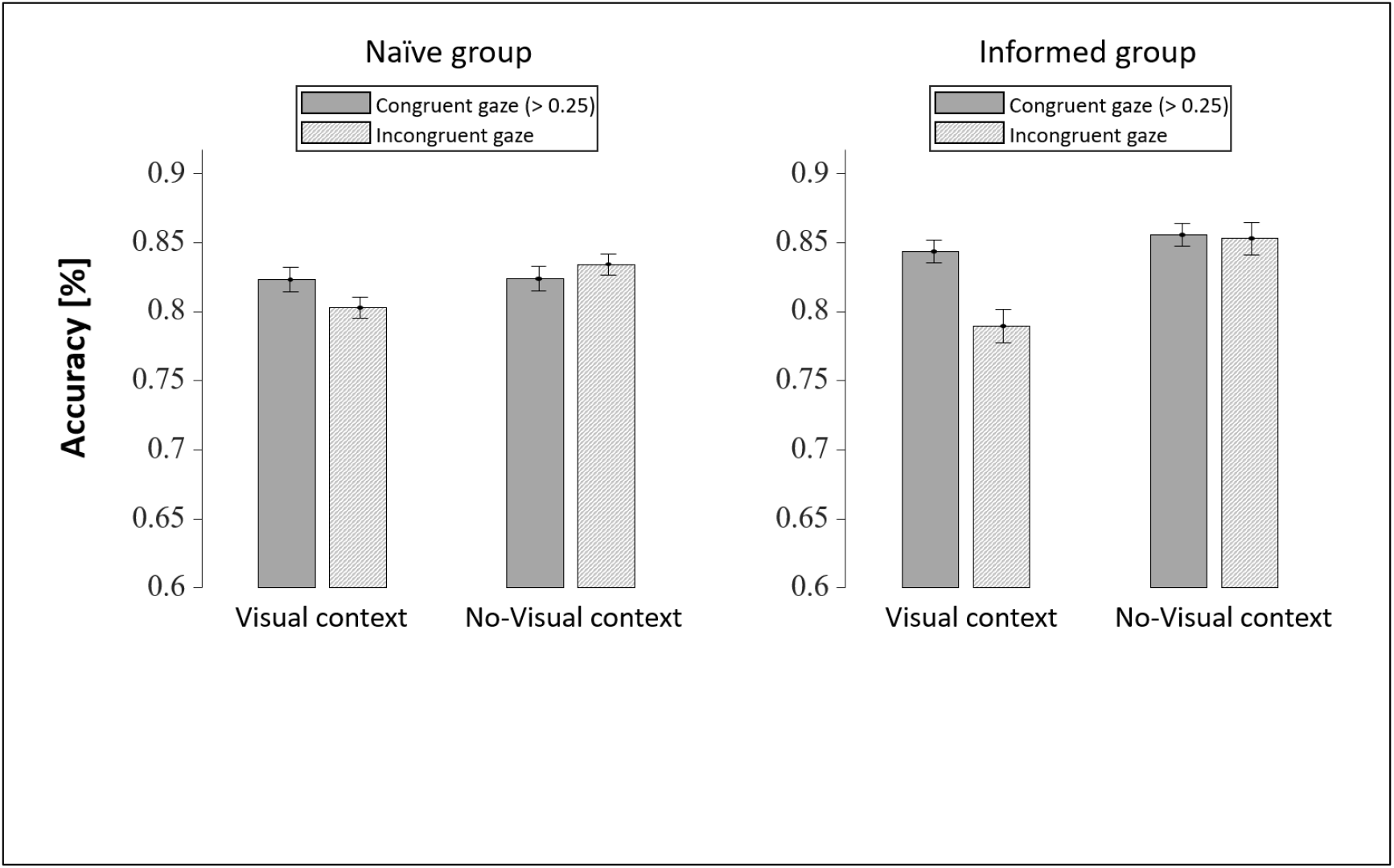
Average accuracy rates for trials of congruent vs. incongruent gaze patterns. Accuracy rates were higher for congruent relative to incongruent gaze trials over both groups in the visual context condition, but not in the no-visual context condition.

The orthogonal simple effects were also analyzed to examine the differences between the two context conditions, separated for congruent gaze and incongruent gaze trials. We found that for the congruent gaze trials, there was no effect for context, meaning that, for those trials, accuracy was similar in the visual context condition and in the no-visual context condition (t(141.822)=0.320, p=0.749). However, in the incongruent gaze trials, accuracy was surprisingly higher for no-visual context compared with the visual context condition (t(141.822)=3.503, p<0.001). Taken together, these results indicate that, rather than *gaining* from a congruent gaze itself, memory was *impaired* by an incongruent gaze at the presence of visual context. This suggests that the presence of irrelevant visual input, such that provided by an incongruent gaze, impairs memory (see Figure 3).

We found no evidence for a main effect of Group (F(1)=0.703, p=0.404), nor was there a significant interaction between Group and any of the other two conditions (interaction between Group and Context: F(1)=0.900, p=0.346; interaction between Group and Gaze congruency: F(1)=2.109, p=0.151) or both (three-way interaction between Group, Context and Gaze congruency: F(1)=0.533, p=0.468).

Despite the lack of interaction, we have examined the effect of congruity separately for each group and visual context condition. We tested the simple effects of the interaction between Gaze congruency and Context, now separately for each group. First, we tested each of the Context conditions, to test for differences in accuracy between congruent and incongruent gaze. For the naïve group, we found no evidence for a difference in accuracy between congruent and incongruent gaze for each of the Context conditions (visual context: t(147.841)=1.243, p=0.216; no-visual context: t(147.841)=0.636, p=0.526). For the informed group, we found higher accuracy for congruent vs. incongruent gaze in the visual context condition (t(147.841)=3.295, p=0.001) but not in the no-visual context condition (t(147.841)=0.065, p=0.948; see Figure 3).

We further inspected the orthogonal simple effects, to check for differences in the Context condition according to Gaze congruency in each group. For the naïve group, we found a marginally significant effect of higher accuracy for the visual relative to the no-visual context condition in the congruent trials (t(141.822)=1.675, p=0.0.096), and no evidence for such an affect in the congruent trials (t(141.822)=0.030, p=0.976). Similarly, for the informed group we found significantly higher accuracy rates for the visual condition than the no-visual condition in the incongruent gaze trials (t(141.822)=3.239, p=0.001), but no evidence for such an effect in the congruent gaze trials (t(141.822)=0.413, p=0. 680).

## Discussion

In this study, we have examined the relation between gaze behavior during memory retrieval and the presence of visual context. Consistently with previous studies, our findings show that when remembering a visual object, people tend to look at the location where that object was presented during encoding. This behavior occurred both when the visual context was present and when it was occluded. Most importantly, the effect of this behavior on memory performance depended on the presence of visual input. Specifically, we found that gazing at the location where the memorized item was previously presented was associated with higher memory performance only when the visual context was present. When the visual context was occluded, there was no beneficial effect for this gazing behavior. These findings challenge other interpretations of the ‘looking at nothing’ effect as reflecting the role of eye movement as a retrieval cue. Here we suggest that it is not the eye movements themselves, as motor actions, that serve the role of retrieval cues. Rather, it is the *consequences* of eye movements – the visual stimulation that is *caused* by them – that modulates memory performance.

### Looking at congruent locations in the absence of visual context

The findings show that participants spent more time looking at the congruent quadrants, even when visual information was unavailable. This is consistent with previous studies that suggested that during recall, people tend to look at locations associated with the remembered items (Altmann, 2004; Brandt & Stark, 1997; Henderson & Castelhano, 2005; Hoover & Richardson, 2008).

Our results indicated that participants looked towards congruent locations, even when this behavior had no beneficial effect on memory performance. It is an open question what is the purpose of this behavior, if there is any. One possible explanation is that activation of a memorized representation of an object leads to an activation of the representation of its spatial location, which, in turn, triggers an eye movement toward that location. This hypothesis is in line with studies suggesting that the cognitive system utilizes spatial indices in memory tasks, even if the spatial location is task-irrelevant (Richardson & Spivey, 2000). In the current experiment, items were presented by categories, each appearing in a separate quadrant. When participants heard a statement mentioning one of the objects, this could have activated the memory representation of the category’s quadrant, even if there was no recollection of specific details regarding the object itself. The activation of the quadrant location may have caused the eyes to shift towards this quadrant, regardless of a specific memory of the individual item. This could explain why the eyes tend to shift to congruent locations even in the absence of visual context and even when this behavior provides no benefit.

### The association between gaze congruency and accuracy

The findings showed that the difference in accuracy found between the four trial types (congruent/incongruent gaze with/without visual context), stemmed mainly from lower performance in trials in which the gaze was incongruent and visual context was present. We found no difference in memory performance between congruent trials performed with visual context and trials performed when there was no-visual context, regardless of their congruity. This was a surprising finding as most previous literature in the field refers to the “looking at nothing” behavior (termed here ‘congruent’) as a method for *enhancing* memory (e.g., Johansson & Johansson, 2014; Scholz et al., 2016), but the present results suggest that the opposite is true –looking at incongruent locations *impairs* it. This leads to the hypothesis that the purpose of performing eye movements during retrieval is not to collect visual information which could be used as a retrieval cue, but rather to *avoid* collecting distracting information, which has the potential to mislead and decrease performance.

In this study, retrieval conditions differ from one another with one of them preserving the original visual context provided during encoding, and another altering the visual context of the near-surroundings, leaving it almost blank. This absence of competing stimuli could explain why performance was similar in congruent trials with visual context and in trials in which the visual context was removed, regardless of congruity. According to one possible interpretation, when there is no distracting visual input, participants can more easily preserve a mental representation of the original scene and recreate eye movements accordingly. In our no-visual context condition, since there were no competing stimuli and no distracting visual information, performing incongruent gaze shifts did not interfere with memory performance. To examine this question in the future would require alternating the visual scene while preserving its visual saliency, rather than eliminating most of the visual stimulation as was done in the present study.

### The effect of explicit instructions on gaze behavior

The experiment was conducted with two separate groups of participants: one informed of the benefit of gazing at congruent locations and encouraged to do so, and another not informed or encouraged. Our findings reveal that this information and encouragement had a considerable effect on the participants’ behavior, increasing the percentage of dwell time at the congruent quadrant. This indicates that eye movements during retrieval can be voluntarily manipulated top-down by the participant’s explicit intentions and goals. Surprisingly, however, despite the substantial effect of this information on gaze behavior, there was no evidence for an overall improvement in memory performance in the informed group compared to the naïve group.

This is consistent with findings suggesting that intentional memory instructions do not always improve memory but could instead enhance attentional processes during the task (Varakin & Hale, 2014). It is possible that the intentional instruction to encode location and to use it during retrieval may have caused participants to focus attention more on their gaze location during retrieval rather than on the correct retrieval, leading them to less gain in their memory performance. Further research is needed to determine the influence of explicit eye movement strategies on performance in memory tasks.

## Conclusion

In this study, we show that the link between congruent eye movements during memory retrieval and higher memory performance, is dependent on the availability of the visual context. Although such congruent eye movements occur even when the visual context is occluded, they do not lead to higher memory performance. Moreover, the evidence opposes the intuitive hypothesis that gazing at congruent locations would enhance memory performance; in fact, we find that gazing at incongruent locations impairs it. Together, these findings challenge the idea that eye movements serve as a retrieval cue. They instead support a visual-sensory interpretation of previous findings – that receiving irrelevant visual input hinders memory retrieval.

## Acknowledgements

This research was funded by the Israel Science Foundation, grant 1960/19 to S.Y-G and by the Ariane de Rothschild Women Doctoral scholarship to K.T.

